# X-Linked Cancer-Associated Polypeptide (XCP) from *lncRNA1456* Cooperates with PHF8 to Regulate Gene Expression and Cellular Pathways in Breast Cancer

**DOI:** 10.1101/2025.03.21.644649

**Authors:** Shrikanth S. Gadad, Cristel V. Camacho, Xuan Gong, Micah Thornton, Venkat S. Malladi, Anusha Nagari, Aishwarya Sundaresan, Tulip Nandu, Sneh Koul, Yan Peng, W. Lee Kraus

**Author notes:** Current address: Center of Emphasis in Cancer, Paul L. Foster School of Medicine, Department of Biomedical Sciences, Texas Tech University Health Sciences Center, El Paso, TX 79905, USA. Current address: Department of Bone Marrow Transplantation and Cellular Therapy, St. Jude Children’s Research Hospital, Memphis, TN 38105. These authors contributed equally to this work (Shrikanth S. Gadad, Cristel V. Camacho). Lead Contact / Address correspondence to: W. Lee Kraus, Ph.D., Cecil H. and Ida Green Center for Reproductive Biology Sciences The University of Texas Southwestern Medical Center at Dallas 5323 Harry Hines Boulevard Dallas, TX 75390-8511.

## Abstract

Recent studies have demonstrated that a subset of long “noncoding” RNAs (lncRNAs) produce functional polypeptides and proteins. In this study, we discovered a 132 amino acid protein in human breast cancer cells named XCP (<u>X</u>-linked <u>C</u>ancer-associated <u>P</u>olypeptide), which is encoded by *lncRNA1456* (a.k.a. *RHOXF1P3*), a transcript previously thought to be noncoding. *lncRNA1456* is a pancreas– and testis-specific RNA whose gene is located on chromosome X. We found that the expression of *lncRNA1456* and XCP are highly upregulated in the luminal A, luminal B, and HER2 molecular subtypes of breast cancer. XCP modulates both estrogen-dependent and estrogen-independent growth of breast cancer cells by regulating cancer pathways, as shown in cell and xenograft models. XCP shares some homology with homeodomain-containing proteins and interacts with the histone demethylase plant homeodomain finger protein 8 (PHF8), which is also encoded by an X-linked gene. Mechanistically, XCP stimulates the histone demethylase activity of PHF8 to regulate gene expression in breast cancer cells. These findings identify XCP as a coregulator of transcription and emphasize the need to interrogate the potential functional roles of open reading frames originating from noncoding RNAs.

**Statement of Significange:** XCP, a polypeptide encoded by an X-linked lncRNA, regulates gene expression in breast cancer cells. XCP is a chromatin-associated protein that interacts with the histone demethylase PHF8 and modulates its demethylase activity to regulate gene expression. XCP drives cancer-specific phenotypes and serves as a potential biomarker and/or target for therapeutic intervention.

## Introduction

Recent advancements in genomics and deep sequencing technologies have uncovered that a significant fraction of the noncoding mammalian genome is transcribed, leading to the discovery of many long noncoding RNAs (lncRNAs) [1]. While lncRNAs are typically classified as noncoding based on computational approaches (e.g., codon-substitution frequency; CSF) [2], this may not be ideal for some highly evolved RNAs that serve complex biological functions in humans and non-human primates. Additionally, many peptides have gone unnoticed due to technical limitations and assumptions made during the annotation of genome databases [3]. As a result, research has recently focused on identifying functional open reading frames (ORFs) that might be translated into functional novel peptides and proteins derived from transcripts classified as noncoding, particularly lncRNAs.

The remarkable number of expressed lncRNAs, some of which conceivably also have protein-coding potential, have led to a significant gap in our knowledge and understanding of the biological roles of this distinct group of peptides. One of the most interesting features of lncRNAs is their tightly regulated developmental and tissue-specific expression, which is often dysregulated in diseased states, such as cancer [4]. The discovery of lncRNA-derived peptides has the potential to expand our understanding of the function of lncRNAs and their roles in cancer, ultimately uncovering exciting new targets for therapeutic intervention.

Emerging evidence suggests that peptides derived from lncRNAs play a crucial role in physiological processes, as well as in the manifestation of various diseases, including cancer [5–17]. Studies of lncRNAs, such as *long noncoding RNA 152* (*lncRNA152*), which we previously annotated and characterized [18], suggest that they are critical regulators of cancer biology [4]. However, in the studies described herein, we found the presence of potential ORFs in many annotated lncRNAs identified in breast cancer. We focused our analyses on XCP (<u>X</u>-linked <u>C</u>ancer-associated <u>P</u>olypeptide), which is encoded by a pancreas– and testis-specific RNA, *lncRNA1456* (a.k.a. *RHOXF1P3*) [19]. XCP interacts with plant homeodomain finger protein 8 (PHF8) to modulate its demethylase activity and regulate gene expression outcomes in breast cancer. Overall, this study underscores the importance of the novel polypeptide XCP in epigenetic regulation of gene expression and tumor biology.

## Results

### Identification and validation of XCP, a polypeptide encoded by *lncRNA1456*

In this study, we developed a pipeline integrating RNA-sequencing (RNA-seq) and mass spectrometry (MS) data to identify putative functional lncRNA-encoded peptides using a universe of lncRNAs previously discovered and annotated in MCF-7 breast cancer cells [19] (Fig. S1A). Briefly, we curated a peptide database by combining all possible ORFs, starting with methionine from the list of lncRNAs with known proteins from UniProtKB/Swiss-Prot. Further, we perfomed mass spectrometry (MS) on lysates from MCF-7 cells with peptides under 20 kDa, and spectra were analyzed using Proteome Discoverer 2.0 (Thermo Fisher Scientific, Waltham, MA) against the combined peptide database (Fig. S1A). Using this pipeline, we discovered and functionally characterized a new polypeptide encoded by *lncRNA1456*, which we have named XCP (<u>X</u>-linked <u>C</u>ancer-associated <u>P</u>olypeptide) (Fig. S1B).

RNA-seq analysis revealed that *lncRNA1456* is highly transcribed in MCF-7 cells, which is classified as a luminal breast cancer cell line (Fig. 1A). Its transcription is independent of estrogen treatment, and the transcript can be detected in both cytoplasmic and nuclear fractions (Fig. 1A). Analysis of its evolutionary conservation across genomes showed that *lncRNA1456* is conserved in humans and primates only, suggesting this is an evolutionarily young polypeptide (Fig. 1B and Fig. S1B). *LncRNA1456* contains an ORF that is translated into a 132-amino acid peptide with a predicted molecular weight of approximately 14 kDa (Fig. 1C and Fig. S1B). This peptide was detected in MCF-7 whole-cell extracts as a single band at approximately 18 kDa using an antibody raised against recombinant XCP (Fig. 1D). siRNA-mediated knockdown of *lncRNA1456* reduced the levels of detectable XCP polypeptide, confirming antibody specificity (Fig. 1E and Fig. S1C).

**Figure 1.**
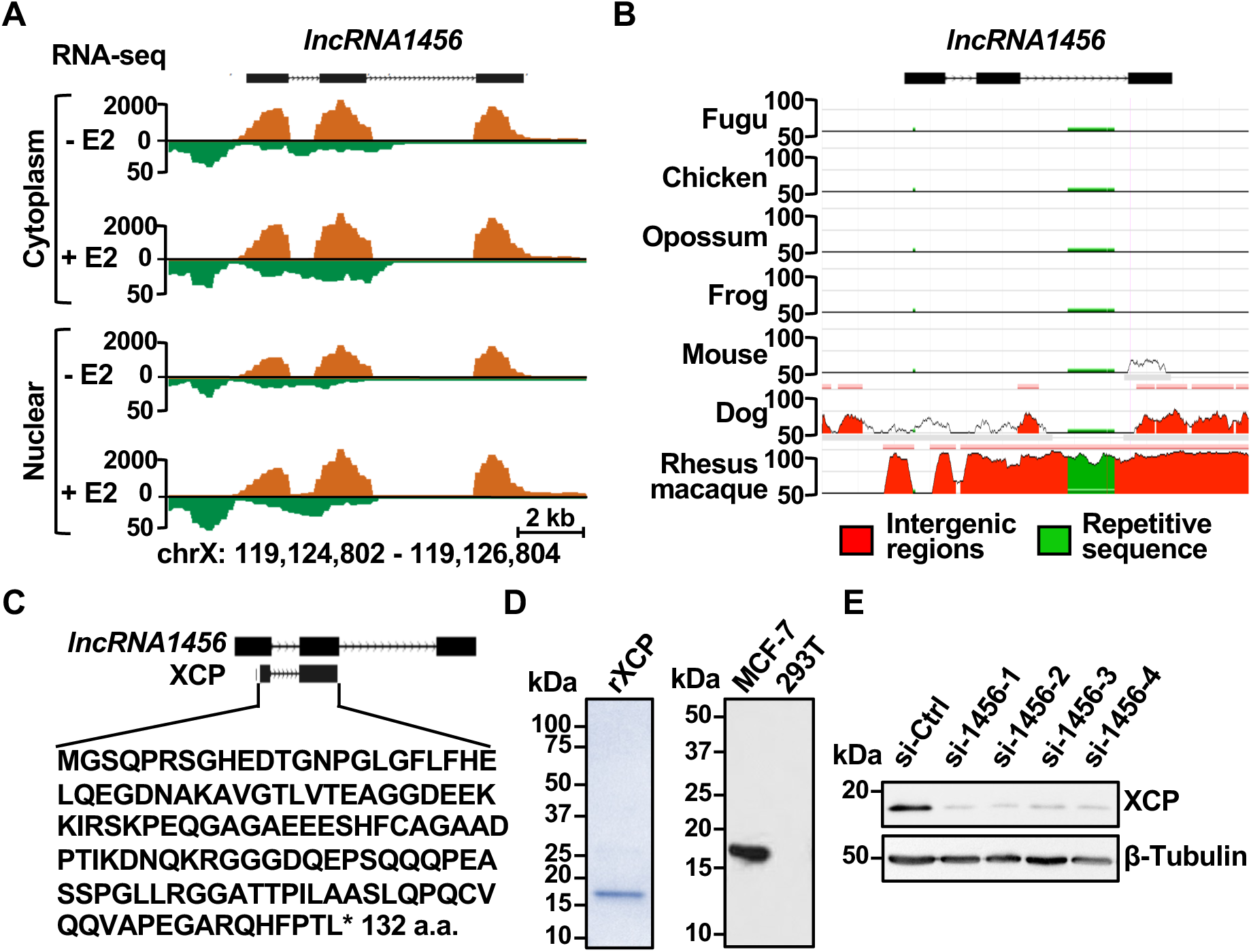
Characterization of *lncRNA1456* and its encoded peptide, XCP. **A)** Genome browser view for the *lncRNA1456* locus from fractionated RNA-seq data. **B)** ECR browser tracks showing conservation of *lncRNA1456* across different species. **C)** Diagram showing the putative *lncRNA1456*-encoded peptide from exons 1 and 2 and amino acid sequence. **D)** Coomassie gel showing recombinant XCP peptide (*left*) and its endogenous detection in MCF-7 cells by Western blot using a derived XCP antibody (*right*). **E)** Western blot showing reduced levels of endogenous XCP peptide upon siRNA-mediated knockdown of *lncRNA1456* RNA in MCF-7 cells. *[See also Figure S1]*

### XCP is highly expressed in luminal subtype breast cancer, promoting tumor cell growth

To rule out the possibility that the XCP peptide is an in vitro artifact, we first determined its RNA levels across normal tissues (GTEx). We found that *lncRNA1456* is transcribed exclusively in the testes and pancreas, which was further confirmed by immunohistochemistry using a collection of normal tissues (Fig. 2A). This expression pattern resembles that of cancer-testis antigens (CTA), which are restricted to adult male germ cells under normal physiological conditions and become aberrantly re-expressed in cancer. In agreement with this, *lncRNA1456* is highly transcribed in primary tumors (T; TCGA) from breast cancer patients but not in normal solid tissue (NT; TCGA) or normal breast tissue (N; GTEx) (Fig. 2B; *left panel*).

**Figure 2.**
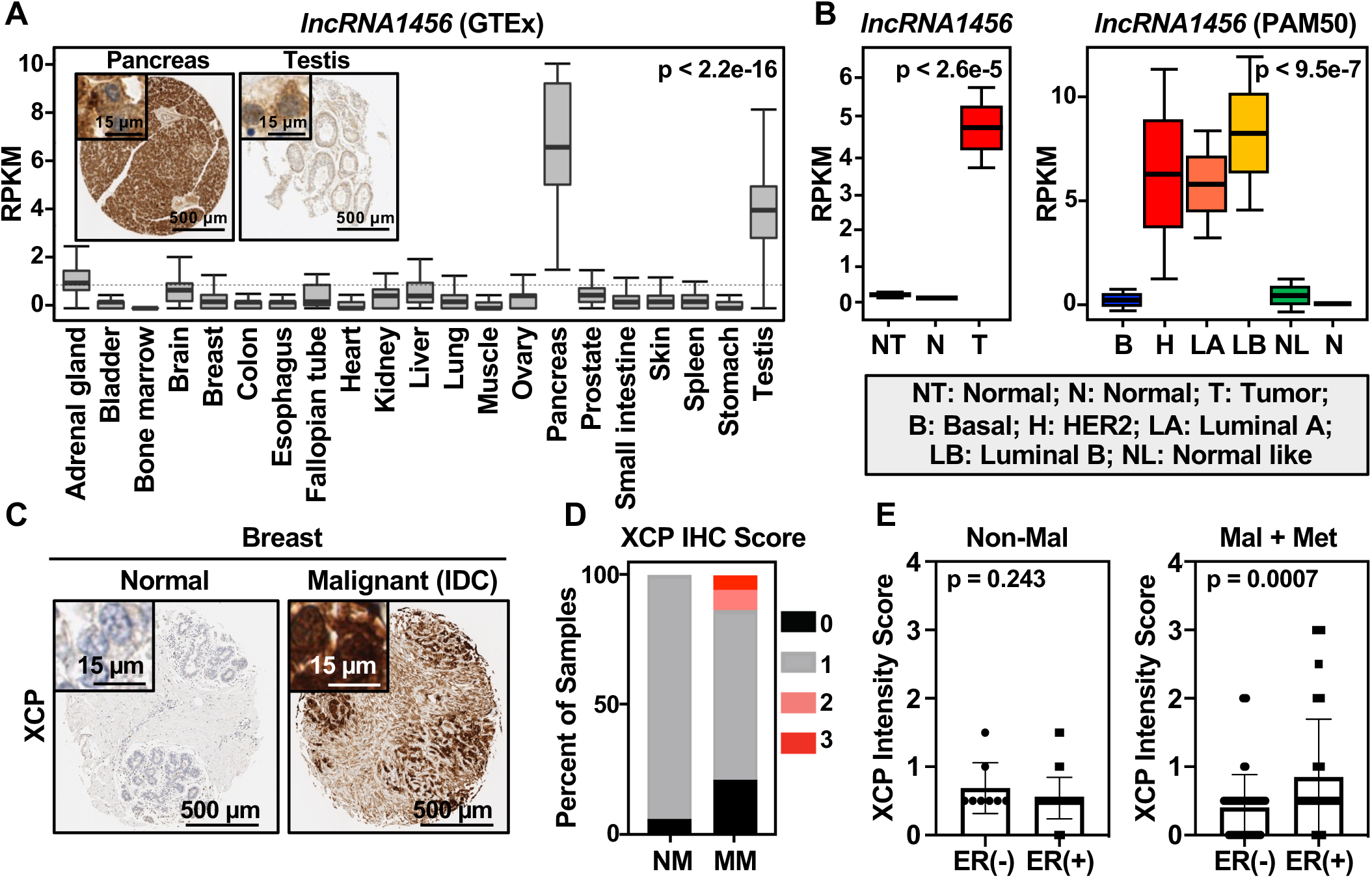
Expression of *lncRNA1456* RNA and XCP peptide in human normal and breast cancer tissues. **A)** GTEx survey of *lncRNA1456* RNA expression across normal human tissues. Observed differences are significant as determined by an ANOVA comparison of the means (p-value < 2.2 e-16). Insets show immunohistochemical staining of pancreas and testis using antibody against XCP. **B)** Graph showing *lncRNA1456* RNA expression in normal solid tissue (NT; TCGA), normal breast tissue (N; GTEx) and malignant primary breast tumors (T; TCGA) (*left*) and *lncRNA1456* RNA expression in malignant primary breast tumors (TCGA) stratified into subtypes (B: basal; H: HER2; LA/LB: luminal A/B; NL: normal like) using PAM50 gene set analysis compared to normal breast tissue (N; GTEx) (*right*). Observed differences are significant as determined by an ANOVA comparison of the means. **C)** Representative immunohistochemical staining for XCP peptide in normal versus malignant (IDC) breast tissue. **D)** Frequency of XCP peptide expression across non-malignant (NM) and malignant plus metastatic (MM) breast tissues based on IHC staining intensity scores. **E)** Expression scores based on IHC of XCP in non-malignant (*left*) and malignant plus metastatic (*right*) breast cancer samples, stratified by ER status. Each bar represents the mean ± SEM; For Non-Mal, ER(–) n=8 and ER(+) n=36; For Mal+Met, ER(–) n=53 and ER(+) n=89. Significance was calculated using unpaired t-test.

Analysis of XCP peptide expression using a commercial breast cancer tissue microarray indicates that XCP is expressed at high levels only in malignant and metastatic (MM) breast cancer tissues but not in non-malignant (NM) tissue (Fig. 2, C, and D); within malignant and metastatic tissues, XCP was found to be significantly more highly expressed in estrogen receptor-positive (ER+) samples (Fig. 2E). Interestingly, and in agreement with this observation, although the overall frequency of XCP-positive samples was low, *lncRNA1456* was found to be upregulated in HER2, luminal A, and luminal B molecular subtypes of breast cancer (TCGA; Fig. 2B; *right panel*), suggesting a context-specific biological role. To explore this further, we ectopically expressed doxycycline (Dox)-inducible XCP-FLAG in MCF-7 and MDA-MB-231 breast cancer cell lines, representing a luminal and basal subtype of breast cancer, respectively, and examined cell growth in a xenograft model. Ectopic expression of XCP-FLAG in MCF-7 (luminal) cells significantly promoted tumor cell growth (Fig. 3, A and C). In contrast, an inhibitory effect was observed when ectopically expressed in MDA-MB-231 (basal) cells (Fig. 3, B and D), indicating that XCP is playing a dual (oncogenic and tumor suppressive) role in a context-dependent manner.

**Figure 3.**
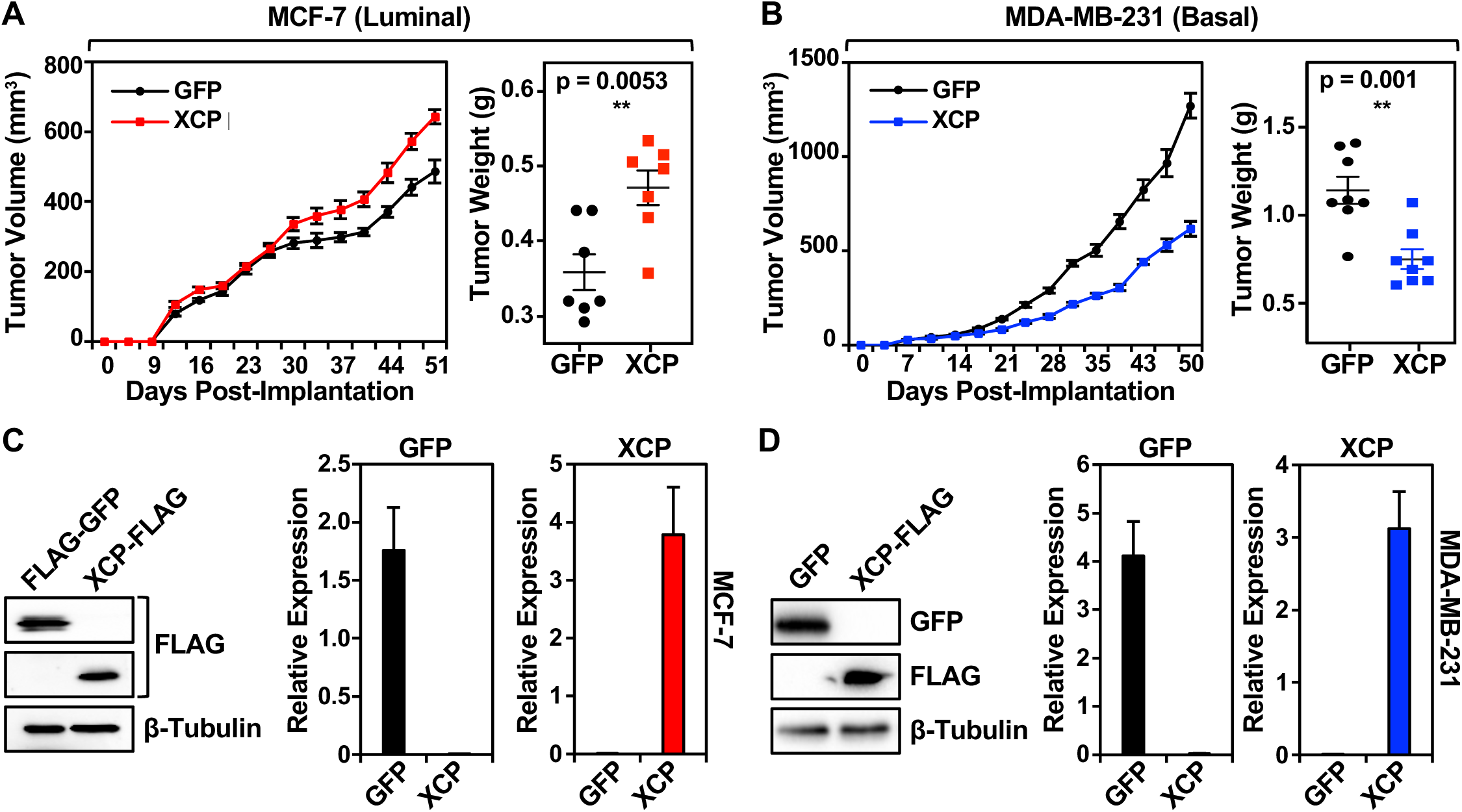
XCP modulates tumor growth in a context-dependent manner. **A and B)** Xenograft assays showing subcutaneous tumor growth of MCF-7 (A) or MDA-MB-231 (B) breast cancer cells ectopically expressing dox-inducible FLAG-GFP or XCP-FLAG (*left*). Tumor weight at end of experiment is shown (*right*). Animals injected with MCF-7 cells had E2-pellet implantations at the back of the neck to promote growth of MCF-7 cells in vivo. All animals were fed doxycycline in their chow. Each point represents the mean ± SEM. For MCF-7, GFP n=7 and XCP n=7; For MDA-MB-231, GFP n=8 and XCP n=8. Significance was calculated using unpaired t-test. **C and D)** Expression of FLAG-GFP or XCP-FLAG in tumor tissue was validated from MCF-7 (**C**) or MDA-MB-231 (D) xenografts tumors by Western blot (*left*) and RT-qPCR (*right*). Each bar represents the mean + SEM. For MCF-7, GFP n=5 and XCP n=5; For MDA-MB-231, GFP n=8 and XCP n=8.

### XCP modulates tumor growth and regulates cellular pathways in a context-dependent manner

To better understand the biological consequences of XCP expression, we performed RNA-seq on MCF-7 and MDA-MB-231-derived xenograft tumors. Ectopic expression of XCP-FLAG resulted in differential gene expression in each cell line (Fig. 4, A and B). Gene set enrichment analysis (GSEA) showed enriched biological pathways that are regulated in the context of MCF-7 (luminal) or MDA-MB-231 (basal) cells. Interestingly, and consistent with the opposing effects observed in tumor cell growth, the pathways that were negatively enriched in MCF-7 (such as EMT, myogenesis, angiogenesis) were positively enriched in MDA-MB-231 cells. In contrast, pathways that were positively enriched in MCF-7 cells (such as G2/M checkpoint, E2F targets, and Myc targets) were negatively regulated in MDA-MB-231 cells (Fig. 4, A and B). Interestingly, in agreement with the restricted expression of *lncRNA1456* and XCP observed in testes (Fig. 2A), enrichment of the term ‘spermatogenesis’ suggests a biological role for XCP in the normal physiology of the testes.

**Figure 4.**
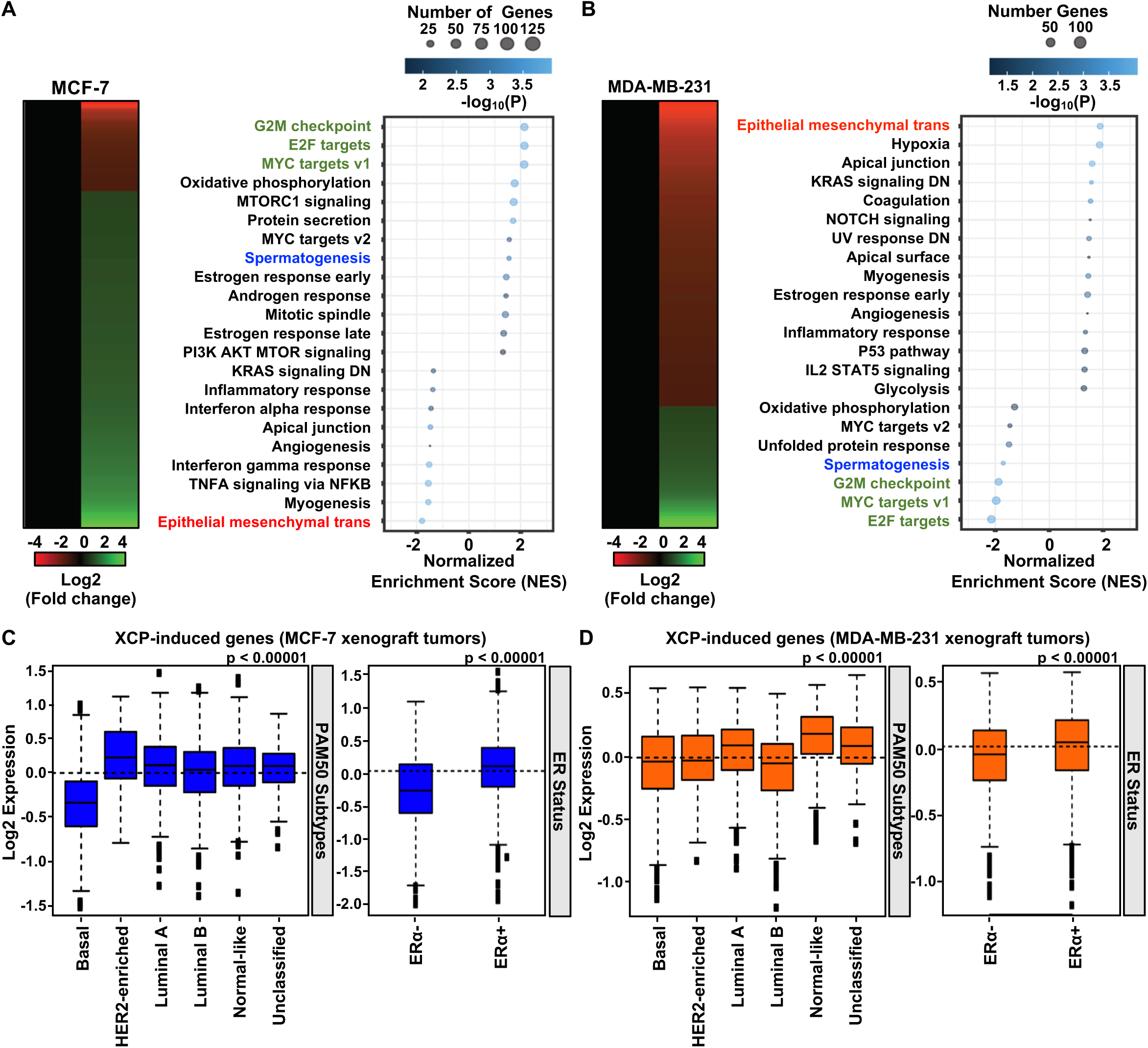
XCP modulates gene expression and regulates cellular pathways in a context-dependent manner. **A and B)** Heatmaps (*left*) and Gene Ontology analysis (*right*) showing XCP-mediated gene expression changes in MCF-7 (A) and MDA-MB-231 (B) xenograft tumors. **C and D)** Box plots showing gene set analysis of XCP-induced genes and their expression levels stratified into breast cancer subtypes using PAM50 gene set analysis (*left*) and stratified by ER status (*right*) in MCF-7 (C) and MDA-MB-231 (D) xenograft tumors. Observed differences are significant as determined by an ANOVA comparison of the means (P value < 0.00001).

We determined the expression of the XCP-induced gene set from MCF-7 xenograft tumors in breast tumor samples categorized by molecular subtype (PAM50) (Fig. 4C). Expression of the gene set was significantly upregulated in the non-basal subtypes of breast cancer (HER2+, Luminal A, Luminal B, etc.), also in tumors positive for ER-alpha (ER11) expression, corroborating the pro-tumorigenic effect of XCP in MCF-7 xenograft tumor growth (Fig. 3A). Intriguingly, the expression of XCP-induced genes from MDA-MB-231 xenograft tumors was also elevated in the luminal subtype of patient breast tumors including ER11+ tumors (Fig. 4D), suggesting a role for XCP in driving the gene expression profiles of luminal subtype tumors.

### XCP interacts with the demethylase PHF8

Immunofluorescent staining of MCF-7 cells demonstrated XCP expression localized to the nucleus (Fig. S2A). A homology search identified homeobox-containing proteins as distantly related protein sequences (Fig. S2B). However, the regulation of gene expression by XCP and a lack of a DNA-binding domain prompted us to speculate that XCP may be acting on chromatin by interacting with a partner protein. To identify possible XCP binding partners and clues about the functions of XCP, we performed MS analysis of XCP-FLAG complexes pulled down from MCF-7 cell lysates (Fig. 5A). Among a number of candidates, MS analysis revealed a clear interaction with the histone demethylase, PHF8 (Table S1). This was confirmed in an in vitro pull down assay using PHF8-FLAG, in which XCP was also detected (Fig. 5B).

**Figure 5.**
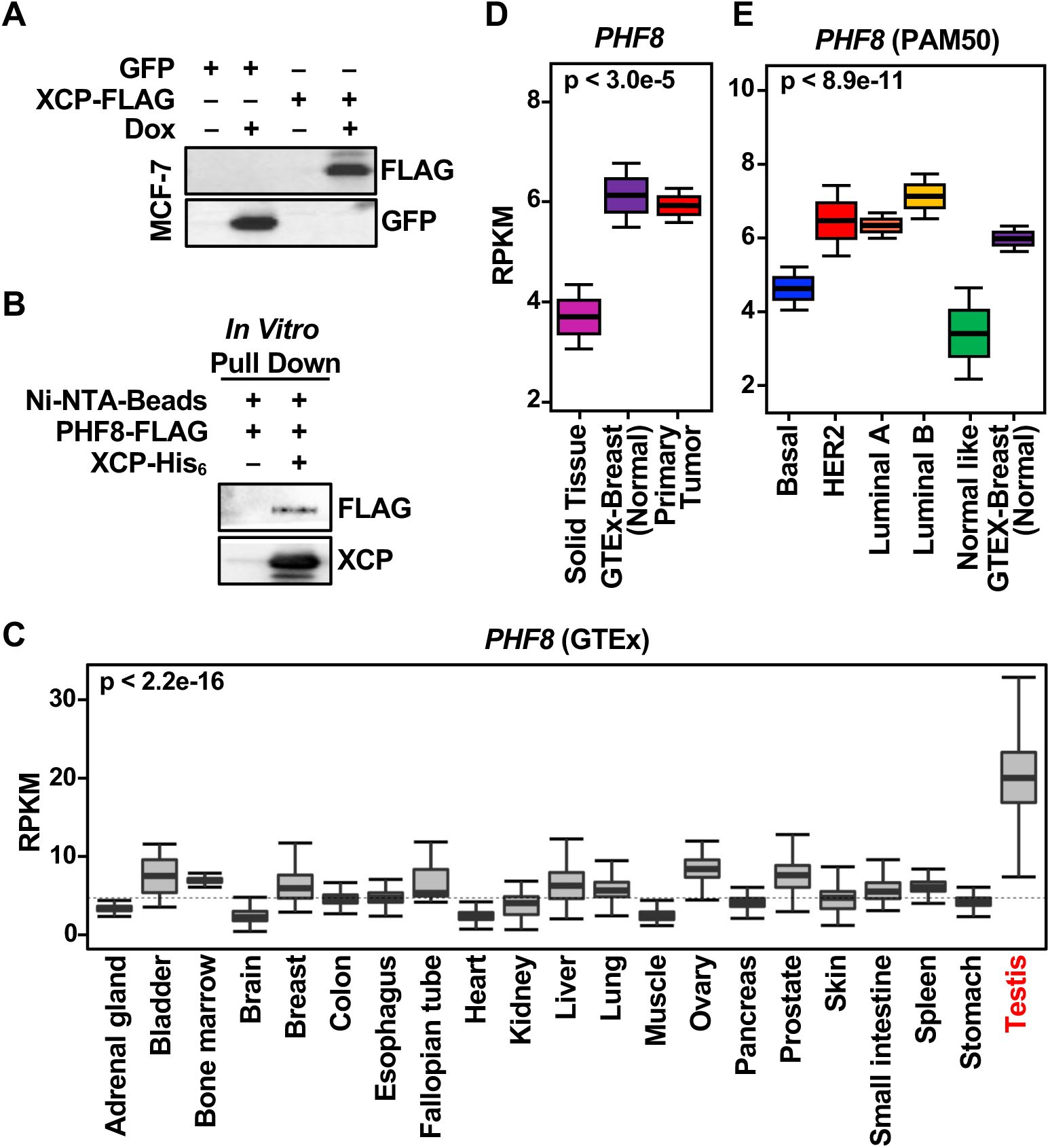
XCP interacts with the demethylase PHF8. **A)** Western blot showing expression of GFP and FLAG-tagged XCP used for pull down from MCF-7 whole cell lysates. **B)** Western blot showing interaction of PHF8 and XCP confirmed in an in vitro pull down assay using recombinant PHF8 and XCP. **C)** GTEx survey of *PHF8* expression across normal human tissues. Observed differences are significant as determined by an ANOVA comparison of the means (p-value < 2.2 e-16). **D)** Graph showing *PHF8* RNA expression in normal solid tissue (TCGA), normal breast tissue (GTEx) and malignant primary breast tumors (TCGA). Observed differences are significant as determined by an ANOVA comparison of the means. **E)** *PHF8* RNA expression in malignant primary breast tumors (TCGA) stratified into subtypes using PAM50 gene set analysis compared to normal breast tissue (GTEx). Observed differences are significant as determined by an ANOVA comparison of the means. *[See also Figure S2]*

Further rationale in support for the selection of PHF8 as an XCP-interacting candidate to study further was the similarities its expression shared with *lncRNA1456* expression: (1) location of the gene to the X chromosome; (2) high expression restricted to the testes in normal tissues (GTEx; Fig. 5C); (3) higher expression levels in normal breast and primary tumor tissues compared to normal solid tissues (Fig. 5D); and (4) higher expression in luminal A, luminal B, and HER2 compared to the basal molecular subtype of breast cancer (Fig. 5E). Collectively, this evidence supports the conclusion that XCP and PHF8 are co-expressed in the same tissues and can interact to regulate biological outcomes.

### XCP regulates gene expression by modulating the binding of PHF8 to chromatin

Previous studies have demonstrated that PHF8 is a histone methyltransferase that acts as a transcriptional coregulator by removing repressive H3K9me2 and H3K9me1 modifications from chomatin [20–24]. To further investigate the function of the interactions between XCP and PHF8, we performed RNA-seq using MCF-7 cells ectopically expressing FLAG-GFP (as a control) or XCP-FLAG, with siRNA-mediated knockdown of either *lncRNA1456* RNA or *PHF8* mRNA, aimed at elevating or reducing XCP protein levels (Fig. S3, A and B). Using the overlap of two different siRNAs for each target, we identified 1,449 and 1,070 genes commonly regulated by *lncRNA1456* or *PHF8* knockdown, respectively (1.5 fold change cutoff). Among these, 239 genes were commonly regulated by both *lncRNA1456* and *PHF8* knockdown. Heat maps using these 239 genes illustrate that a subset of genes differentially regulated upon *lncRNA1456* knockdown can be rescued upon ectopic expression of XCP-FLAG (Fig. 6, A-C; lane 2 versus lane 3). However, rescue of the genes differentially regulated upon *PHF8* knockdown was attenuated upon ectopic expression of XCP-FLAG (Fig. 6, A-C; lane 5 versus lane 6). This suggests that PHF8 is necessary for XCP-mediated gene regulation.

**Figure 6.**
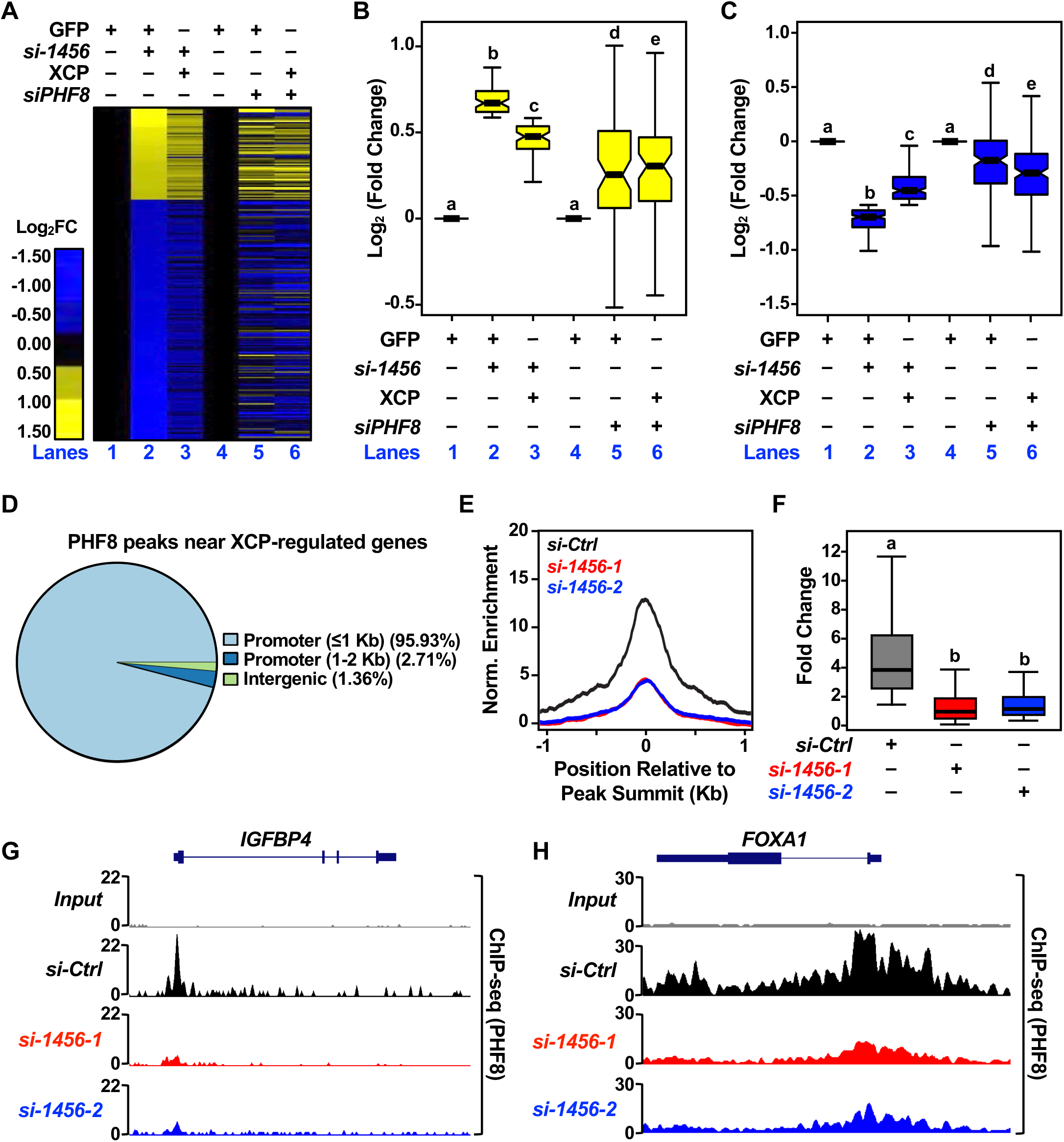
XCP modulates gene expression through its interaction with the demethylase PHF8. **A)** Heatmap from RNA-seq data showing the regulation of 239 genes upon siRNA-mediated knockdown of *lncRNA1456* RNA or *PHF8* mRNA and co-expression of XCP-FLAG. **B and C)** Box plots quantifying the upregulated (B) and downregulated (C) gene expression changes observed in heatmap of 239 XCP-regulated genes (A). Each bar represents the mean ± SEM; Bars marked with different letters are significantly different from each other, Wilcox rank sum test. **D)** Pie chart showing the distribution of PHF8 peaks at the 239 XCP-regulated genes. **E and F)** Metagene plot (E) and box plot (F) showing the reduced enrichment of PHF8 binding upon siRNA-mediated knockdown of l*ncRNA1456*. Each bar represents the mean ± SEM; Bars marked with different letters are significantly different from each other, Wilcox rank sum test. **G and H)** Browser tracks showing reduced PHF8 binding near promoter of *IGFBP4* and *FOXA1* genes. *[See also Figure S3]*

Next, we performed ChIP-seq analysis to assess PHF8 binding across the genome following knockdown of *lncRNA1456* to reduce the levels of XCP. We observe that PHF8 binds primarily near promoters, both globally (Fig. S3C) and near the 239 genes that are regulated by XCP (Fig. 6D). For XCP-regulated genes, PHF8 peaks are enriched within 200 kb of the transcription start sites (TSSs) (Fig. S3D). Interestingly, we detected a dramatic loss of PHF8 binding upon reduction in XCP levels, again both globally and near XCP-regulated genes (Fig. 6, E and F, Fig. S3E). For example, the *IGFBP4* and *FOXA1* genes, both found to be modulated by XCP expression, show reduced PHF8 binding near promoters in response to reduced levels of XCP (Fig. 6, G and H). These results indicate that XCP directly controls the chromatin binding of PHF8 to regulate gene expression.

### XCP modulates the demethylase activity of PHF8

To understand how XCP modulates PHF8 activity, we carried out in vitro demethylase reactions using recombinant PHF8-FLAG, XCP-His_6_, and H3K9me2 peptide (Fig. 7A; *left panel*). We observed that PHF8 alone has a limited ability to demethylate the H3K9me2 peptide, as determined by dot blotting (Fig. 7A; *right panel*, lane 2). However, in the presence of XCP, PHF8 was able to robustly demethylate the H3K9me2 peptide (lane 4), and its activity was further enhanced with increasing amounts of XCP (lane 5). XCP alone did not alter the methylation of the H3K9me2 peptide (Fig. 7A; *right panel*, lane 3).

**Figure 7.**
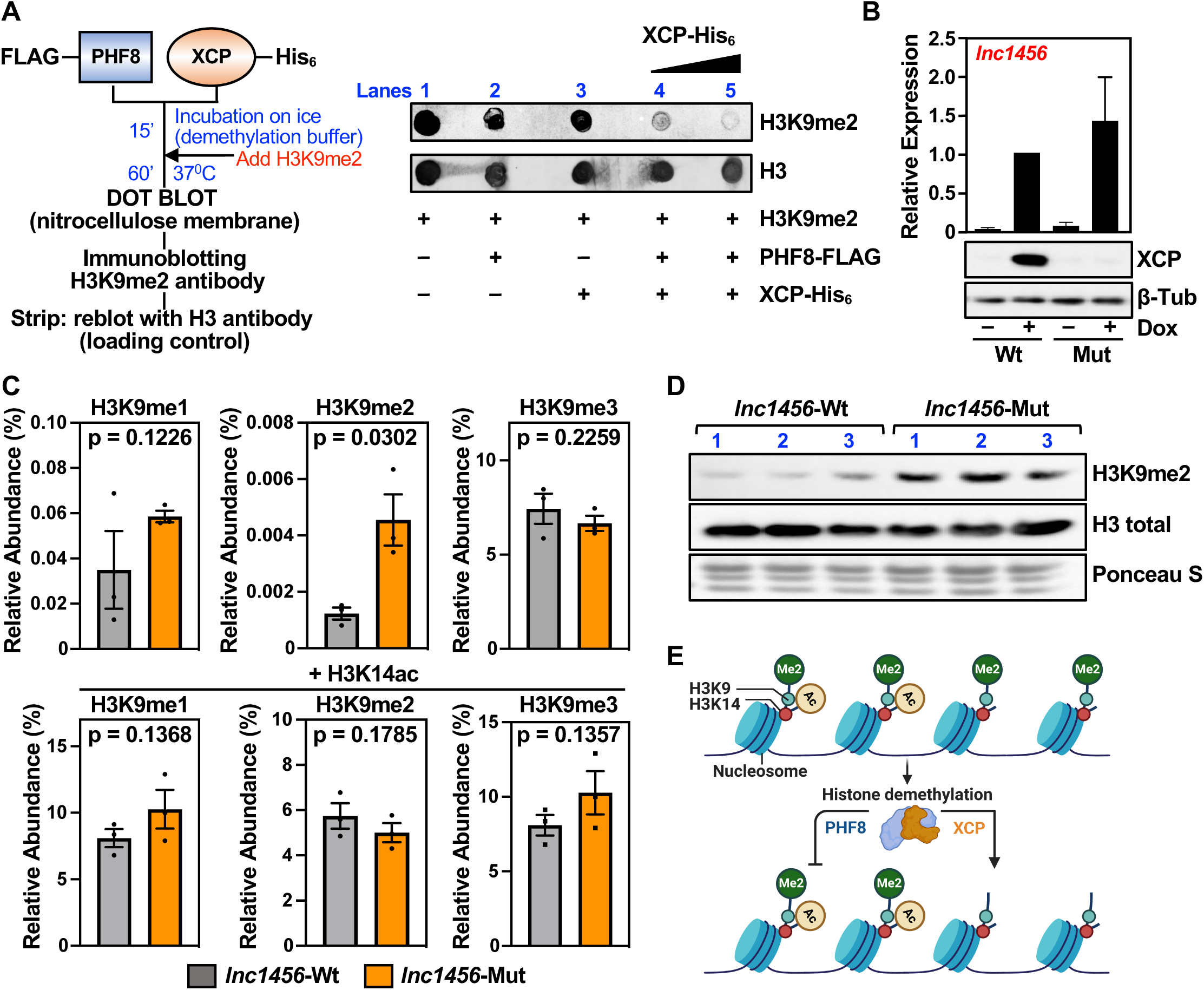
XCP directly modulates the demethylase activity of PHF8. **A)** Diagram showing the flow of in vitro demethylase reactions using recombinant PHF8-FLAG, His_6_-XCP and H3K9me2 peptide (*left*). Dot bot showing the demethylase activity of PHF8 on H3K9me2 peptide in the presence or absence of XCP (*right*). **B)** RT-qPCR (*top*) and Western blot (*bottom*) showing dox-inducible ectopic expression of full length *lncRNA1456* in its wild-type form, which expresses the XCP peptide, or an ATG mutant version which abolishes XCP peptide production. Each bar represents the mean + SEM; n= 3. **C)** Bar graphs showing global changes in histone post-translational modifications assayed mass spectrometry of MCF-7 histones. Each bar represents the mean ± SEM; n=3. Significance calculated by 1-sided Welch’s t test. **D)** Western blot analysis of MCF-7 histone preparation used for mass spectrometry analysis (C). **E)** Diagram summarizing model: XCP, encoded by *lncRNA1456*, interacts with PHF8, a histone demethylase, and directly regulates its activity to modulate gene expression.

To examine the effect of XCP on PHF8-mediated demethylation globally in cells, we examined post-translational modification of histones by mass spectrometry. We isolated histones from MCF-7 cells subjected to siRNA-mediated knockdown of *lncRNA1456* and ectopic co-expression of full-length *lncRNA1456*, which expresses XCP, or an XCP start codon (ATG) mutant that abolishes XCP production (Fig. 7B). We assayed methylation marks that are known to be the preferred targets of PHF8 by mass spectrometry. We observed a significant reduction in H3K9me2 levels with expression of wild-type *lncRNA1456* (XCP expressed) compared to the ATG mutant (XCP absent) (Fig. 7C; *top panels*). This was validated by Western blotting for H3K9me2 on isolated histones, strengthening our observations (Fig. 7D). Interestingly, acetylation at the neighboring H3K14 residue abrogates the effect on H3K9 dimethylation in wild-type versus mutant *lncRNA1456* expressing cells (Fig. 7D*; bottom panels*). Overall, these results provide evidence for the global modulation of PHF8 demethylase activity by XCP, resulting in the transcriptional regulation of gene expression in breast cancer cells (Fig. 7E).

## Discussion

Expansion of genome-wide transcriptome analysis has revealed that mammalian genomes are pervasively transcribed, and the ‘noncoding RNA’ genome has garnered significant attention in recent years. LncRNAs are an interesting species of noncoding RNAs exhibiting a high level of cell type and developmental specificity; their expression often becomes dysregulated in disease states. For example, cancers carry a heavy mutational load in the noncoding genome, which could profoundly affect the expression of lncRNAs and, hence, cellular biology [25, 26]. Recently, it has become clear that some lncRNAs, which are sometimes misannotated as non-protein-coding, may actually contain translatable open reading frames (ORFs). Many lncRNAs that have been categorized as non-protein coding are often misclassified as such due to arbitrary cutoff points for ORF sizes set by computational algorithms [27]. As a result, many potential ORFs were often marked as evolutionarily dispensable and, hence, not translatable. These ORFs can vary in size, producing anything on the order of 100 amino acids. In the last five to ten years, there is significant evidence from proteomics and ribo-seq profiling showing that these ORFs, many encoded within this ‘noncoding RNA’ genome, are abundant, evolutionarily young, and biologically functional [9, 28–31]. The lack of characterization of specific cellular roles for these small functional peptides remains a gap in our knowledge.

### XCP is an X-linked cancer-testis antigen

We have identified a lncRNA-encoded peptide, which we have named XCP, and demonstrated its biological relevance in breast cancer cells. The gene encoding XCP, *lncRNA1456*, like many others transcribed in the testis, is located on the X chromosome, and its sequence conservation is primarily observed in humans and non-human primates. Although XCP gene expression is largely restricted to normal tissues such as the testis and pancreas, it can escape regulation in malignant cells from females, likely due to aberrant epigenetic regulation that presumably leads to the reversal of X-inactivation. In fact, the majority of novel transcripts (coding and noncoding) that are frequently identified from cancer cells have restricted expression in the testis and originate from the X chromosome [32]. We propose, based on the aberrant expression of XCP in breast cancer cells, and its encoding on the X chromosome, that XCP can be classified as an X-linked cancer-testis antigen, which could be an interesting new target for antigen-directed immunotherapies in breast cancers.

### XCP has dual, context-dependent functions in gene regulation

Our findings indicate that XCP is upregulated not only at the RNA level, but also at the protein level in breast cancers, particularly in non-aggressive breast cancer (luminal subtype), while it is found at reduced or minimal levels in more aggressive cases (basal subtypes). This is in agreement with our observation that ectopic expression of XCP in MCF-7 cells (luminal or ER+ cells) promotes E2-dependent tumor growth, whereas an inhibition of tumor growth is observed in a MDA-MB-231 (basal or ER–) cell line.

The different effects of XCP in different cellular contexts can be attributed to the specific genes and cellular pathways it regulates in ER+ and ER– cells, as identified through genome-wide expression analyses. Furthermore, the expression patterns of XCP-regulated genes correspond to the breast cancer subtypes. For instance, genes regulated in ER+ breast cancer cells were also expressed in luminal breast cancer subtypes, while genes regulated in triple-negative breast cancer cells were also expressed in basal breast cancer subtypes. This context-dependent regulation may be due to a distinct set of proteins or molecules with which XCP interacts. A similar observation has been made with other proteins specific to epithelial cells, such as FOXA1 or GATA3, which reduce cell growth when overexpressed in basal cell lines [33, 34].

### XCP functions as an epigenomic regulator via intreractions with PHF8

XCP localizes to the nucleus, suggesting a role in the regulation of gene expression, and partners with PHF8, suggesting a role in chromatin regulation. Although homology searches indicated similarities to proteins containing DNA-binding domains, XCP itself does not contain amino acid sequences similar to those domains. This observation led us to hypothesize that XCP interacts with proteins that have DNA-binding capabilities. In this regard, we identified PHF8, which contains a plant homeodomain—a known DNA-binding domain. Notably, similar to XCP, PHF8 expression is restricted to the testis and is upregulated in luminal subtype of breast cancer, while its expression is downregulated in aggressive breast cancers, including basal subtypes. These similarities between XCP and PHF8 strongly indicate their cooperative roles in regulating gene expression.

PHF8 is a histone demodifying enzyme and an epigenomic modulator that has been shown to demethylate H3K9me2, a repressive chromatin modification, thereby activating transcription and regulating gene expression. Through its intreractions with PHF8, XCP functions to support PHF8-driven epigenomic modification and gene expression (Fig. 7E). This is analogous to other coactivators that interact with epigenetic modulators to regulate gene expression; for example, specific coactivators interact with the histone acetyltransferase p300 to enhance acetylation at promoters and promote transcription [35].

Previous studies have shown that coregulators can modulate the enzymatic activity of histone modifiers, such as acetyltransferases and methyltransferases [35]. Specifically, XCP directly interacts with and modulates the methyltransferase activity of PHF8, enhancing demethylation of H3K9me2. Although functional lncRNAs can function independently of protein translation, ectopic expression of a mutant form of *lncRNA1456*, which cannot be translated into XCP, did not retain the same functionality as wild-type *lncRNA1456*. Thus, key functions of *lncRNA1456* require the protein coding of XCP. Given the low expression levels of PHF8 in cells, it is plausible that it relies on interactions with other proteins like XCP, resulting in a synergistic effect that enhances transcriptional regulation efficiency.

Mechanistically, we show that XCP is a chromatin-associated protein that binds to a subset of PHF8-binding regions; this interaction regulates the demethylase activity of PHF8, thereby modulating the gene expression of target genes in breast cancer cells (Fig. 7E). Our study demonstrates how lncRNA-encoded peptides/proteins can drive cancer-specific phenotypes and serve as potential biomarkers and/or targets for therapeutic intervention.

## Acknowledgments

We thank Yasmin M. Vasquez for assistance with ChIP and members of the Kraus lab for their helpful comments and support. We also acknowledge and thank the following UT Southwestern core facilities: the Live Cell Imaging Core for microscopy support (Dr. Katherine Luby Phelps; 1S10OD021684-01), the Next Generation Sequencing Core for deep sequencing services (Dr. Ralf Kittler), the Proteomics Core Facility for mass spectrometry (Dr. Andrew Lemoff), and the Tissue Management Shared Resource at the Simmons Cancer Center (NCI; 5P30CA142543) for immunohistochemical support. S.S.G. is a CPRIT scholar in cancer research and is supported by a first-time faculty recruitment award from the Cancer Prevention and Research Institute of Texas (CPRIT; RR170020). This work was initiated with support from a grant from the Cancer Prevention and Research Institute of Texas (CPRIT; RP190235) and funds from the Cecil H. and Ida Green Center for Reproductive Biology Sciences Endowment to W.L.K. S.S.G. is also supported by grants from the American Cancer Society (RSG-22-170-01-RMC), NIH 1R16GM149497, and CPRIT-TREC (RP230420).

## Author Contributions

S.S.G and W.L.K. conceived this project and oversaw its execution. S.S.G. and C.V.C. designed and performed the experiments. S.S.G. performed all biochemical experiments. X.G. prepared samples for the histone mass spectrometry. M.T. analyzed mass spectrometry data. V.S.M. developed a bioinformatics pipeline to detect ORFs and analyzed TCGA, GTEx, and initial mass-spec data sets. C.V.C. performed the in vivo experiments, analysis of human tissue arrays, and ChIPs. A.N., V.S.M., T.N., and S.K.. analyzed ChIP-seq data. A.N., A.S., and T.N. analyzed RNA-seq data. Y.P. performed pathological review of human tissue arrays. S.S.G. and C.V.C. prepared initial drafts of the figures and text, which were edited and finalized by W.L.K.

## Supplementary Data

Supplemental information includes three figures and one table [See the Supplementary Data file.]

**Figure S1. Identification and characterization of *lncRNA1456* and XCP, a peptide that it encodes**.

**A)** Schematic showing integrated RNA-seq and mass spectrometry pipeline used to identify putative lncRNA-encoded peptides in MCF-7 cells.

**B)** Data box showing key features of *lncRNA1456*.

**C)** Depletion of *lncRNA1456* RNA levels in response to siRNA-mediated knockdown quantified by qPCR in MCF-7 cells. Each bar represents the mean + SEM; n=3.

**Figure S2. Localization and homology of XCP peptide**.

**A)** Immunofluorescence staining of MCF-7 showing nuclear localization of XCP.

**B)** Diagram showing homology of XCP with homeobox-containing proteins ESX1 and RHOXF2B.

**Figure S3. XCP modulates PHF8 DNA binding activity**.

**A)** Western blots showing dual siRNA-mediated knockdown of *lncRNA1456* RNA with ectopic expression of XCP-FLAG (*top*), and siRNA-mediated knockdown of *PHF8* (*bottom*).

**B)** siRNA-mediated knockdowns were also confirmed by RT-qPCR analysis measuring *lncRNA1456* RNA (*top*) and *PHF8* (*bottom*) mRNA levels. Each bar represents the mean + SEM; n=3.

**C)** Pie chart showing the global distribution of PHF8 peaks.

**D)** Bar graph showng the PHF8 peak distance from TSS for the 239 XCP-regulated genes.

**E)** Metagene plot showing the global reduction in PHF8 binding upon siRNA-mediated knockdown of *lncRNA1456*.

**Table S1. PHF8 Identification by Mass Spectrometry Analysis**.

Summary of mass spectrometry analysis identifying PHF8 as an XCP-interacting partner.

## Experimental Procedures

### Cell culture and treatments

MCF-7 cells were kindly provided by Benita S. Katzenellenbogen (University of Illinois, Urbana-Champaign, Champaign, IL) and MDA-MB-231 cells were purchased from the American Type Culture Collection (ATCC). MCF-7 cells were maintained in Minimum Essential Medium Eagle (Sigma-Aldrich, M1018) supplemented with 5% calf serum (Sigma-Aldrich). For experiments involving estrogen treatment, the cells were grown for at least 3 days in phenol red-free Minimum Essential Medium Eagle (Sigma-Aldrich, M3024) supplemented with 5% charcoal-dextran-treated calf serum and then treated with ethanol (vehicle) or 17β-estradiol (E2; 100 nM; Sigma-Aldrich, E8875) as indicated. MDA-MB-231 cells were maintained in RPMI-1640 Medium (Sigma-Aldrich, R8758) supplemented with 10% fetal bovine serum (Sigma-Aldrich). 293T cells were maintained in high glucose Dulbecco’s Modified Eagle’s Medium (Sigma-Aldrich, D5796) supplemented with 10% fetal bovine serum. For doxycycline (Dox; Sigma-Aldrich, D9891) induction, cells were treated with 250 ng/mL of Dox for 24-48 hours. All cell lines were verified for cell type identity using the GenePrint 24 system (Promega, B1870), confirmed as *Mycoplasma*-free every 6 months using the Universal Mycoplasma Detection Kit (ATCC, 30-1012K).

### Antibodies

The custom rabbit polyclonal antiserum against XCP was generated by Pocono Rabbit Farm and Laboratory by using purified recombinant XCP. Other antibodies used were as follows: PHF8 (Bethyl, A301-772A), GFP (Abcam, ab13970), FLAG M2 (Sigma, F3165), DYKDDDDK Tag (Invitrogen, PA1-984B), β-tubulin (Abcam, ab6046), Histone H3K9me2 (Cell Signaling, 9753), Histone H3 (Abcam, ab1791), goat anti-rabbit HRP-conjugated IgG (Pierce, 31460), goat anti-mouse HRP-conjugated IgG (Pierce, 31430), goat anti-chicken HRP-conjugated IgG (Abcam, ab6877), Alexa Fluor 488 goat anti-rabbit IgG (Thermo Fisher Scientific, R37116).

### RNA isolation, and polyA+ RNA-seq

Estrogen-withdrawn MCF-7 cells were treated with ethanol or 100 nM E2 for 3 hours and subjected to cell fractionation into cytoplasmic, nucleoplasmic, and chromatin fractions. Total RNA was isolated from each fraction using the PARIS kit (Ambion, AM1921). Total RNA was isolated from unfractionated MCF-7 cells using the RNeasy kit (Qiagen, 74136) according to the manufacturer’s instructions. The RNA collected was processed for whole genome polyadenylated RNA sequencing (polyA+ RNA-seq).

The total RNA samples were subjected to enrichment of polyA+ RNA as described previously [36]. Briefly, poly(A)+ RNA was enriched using Dynabeads oligo(dT)25 (Invitrogen), heat fragmented, and reverse transcribed using random hexamers in the presence of dNTPs. Second strand cDNA synthesis was performed with dNTPs, but replacing dTTP with dUTP. After end-repair, dA tailing, ligation to adaptors containing barcode sequences, and size selection using AMPure XP beads (Beckman Coulter, A63881), the synthesized second-strand was digested using uracil DNA glycosylase (Enzymatics, Y9180L). A final PCR reaction was performed using KAPA HiFi HotStart Ready Mix (KAPA Biosystems, KK2612). After library quality control assessment using a Bioanalyzer (Agilent), the samples were subjected to 50 bp single-end sequencing using an Illumina NextSeq Sequencing System. At least two biological replicates were sequenced for each cell line with a minimum of ∼65 M raw reads per cell line.

### Rapid Amplification of cDNA Ends (RACE)

Rapid amplification of 5’ and 3’ ends was performed using 5’/3’ RACE Kit, 2^nd^ Generation (Roche, 03-353-621-001), according to the manufacturer’s instructions.

### Computational pipeline for identification of potential lncRNA-derived peptides

#### Database of ORFs

We started with a comprehensive set of annotated lncRNAs from LNCipedia (v4.0) [37] and converted to bed format using UCSC utilities gtfToGenePred and genePredToBed (http://hgdownload.soe.ucsc.edu/admin/exe/). Extraction of sequences for each annotation was achieved by using getfasta in BEDTools (v2.17.0) [38]. All possible open reading frames (ORFs) starting with Methionine were determined using the getorf tool in the European Molecular Biology Open Software Suite [39].

#### Length of ORFs

Histogram representations were used to assess the median length of all ORFs and the longest ORF for each transcript.

#### Protein digest

Maximum theoretical coverage of predicted ORFs and known proteins from UniProt [40] digested by trypsin and chymotrypsin was determined using the MS Proteomics tools library (https://github.com/msproteomicstools/msproteomicstools). Histograms representations were used to assess the median coverage of digestion by either enzyme.

#### Peptide database

We created a comprehensive protein database by combining our database of predicted ORFs with known proteins from UniProtKB/Swiss-Prot [41]. Spectra were analyzed using Proteome Discoverer 2.0 (Thermo Fisher Scientific, Waltham, MA) against the combined protein database.

### Mass spectrometry and analysis

#### Preparation of cell extracts for mass spectrometric analysis

Cells were collected, washed with ice-cold PBS and resuspended in Whole Cell Lysis Buffer [50 mM Tris-HCl pH 7.5, 0.5 M NaCl, 1 mM EDTA, 1% NP-40, 10% Glycerol, and 1x complete protease inhibitor cocktail (Roche, 11697498001)] and incubated for 30 minutes on ice with gentle mixing to lyse the cells and extract the proteins. All lysates were clarified by centrifugation in a microcentrifuge for 5 minutes at 4°C at full speed. The extracts were fractionated on 4-12% gradient SDS-PAGE, followed by Coomassie staining. Bands between 5 to 20 KDa were excised from the gel and subjected to LC-MS/MS.

#### Analysis

The Mass-spectrometry spectra were analyzed using Proteome Discoverer 2.0 (Thermo Fisher Scientific, Waltham, MA) against the combined peptide database described above. Software, scripts and other information about the analyses can be obtained by contacting the corresponding author (W.L.K.).

### Molecular cloning and purification of recombinant XCP

XCP coding sequence was subcloned between the NcoI and XhoI sites of the pET19b vector by PCR-based sub cloning from MCF-7 cDNA library. His_6_ tagged XCP was purified from E. coli cells using Ni-NTA column as described elsewhere [42].

### Preparation of cell extracts and Western blotting

#### Preparation of whole cell lysates

Cells were collected, washed with ice-cold PBS and resuspended in Whole Cell Lysis Buffer [50 mM Tris-HCl pH 7.5, 0.5 M NaCl, 1 mM EDTA, 1% NP-40, 10% Glycerol, and 1x complete protease inhibitor cocktail (Roche, 11697498001)] and incubated for 30 minutes on ice with gentle mixing to lyse the cells and extract the proteins. All lysates were clarified by centrifugation in a microcentrifuge for 5 minutes at 4°C at full speed.

#### Determination of protein concentrations and Western blotting

Protein concentrations were determined using a BCA protein assay (Pierce, 23225). The cell extracts were aliquoted, flash-frozen in liquid N_2_, and stored at −80 °C. Aliquots of the cell extracts were run on polyacrylamide-SDS gels and transferred to nitrocellulose membranes. After blocking with 5% nonfat milk in TBST, the membranes were incubated with the primary antibodies described above in 3% nonfat milk prepared in TBST, followed by anti-rabbit, anti-mouse, or anti-chicken HRP-conjugated IgG. Western blot signals were detected using an ECL detection reagent (Thermo Fisher Scientific, 34077, 34095).

### Functional analyses of lncRNAs in MCF-7 cells

#### siRNA-mediated knockdowns (lncRNA1456 and PHF8) in MCF-7 cells

Transient siRNA-mediated knockdown of *PHF8* or *lncRNA1456* was performed by transfection of commercially available siRNA oligos targeting human PHF8 (siRNA ID # s61133 and s23107; Invitrogen, 4392420), and custom-designed siRNAs for *lncRNA1456* (designed using SciTools RNAi design software from Integrated DNA Technologies). Commercially available control siRNAs (Sigma-Aldrich, MISSION siRNA universal negative control) were also used. Cells were plated at a density of 2 x 10^5^ cells per well in six-well dishes. Transfections were done at a final concentration of 10 nM using Lipofectamine RNAiMAX reagent (Invitrogen, 13778150) according to the manufacturer’s instructions. Forty-eight hours post-transfection, cells were collected for RT-qPCR or RNA-seq.

#### siRNA sequences

We used the following nucleic acid oligonucleotides for targeted knockdowns:

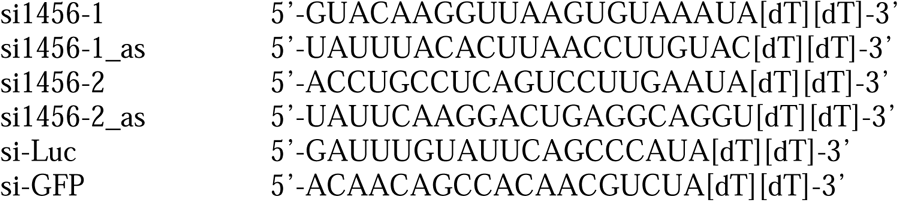

#### Analysis of lncRNA and mRNA expression by RT-qPCR

Total RNA was isolated from MCF-7 or MDA-MB-231 cells using the RNeasy kit (Qiagen, 74136) according to the manufacturer’s instructions. For xenograft tumors, tissue was homogenized in RTL Plus Buffer (Qiagen 74136). RNA was subjected to reverse transcription using Oligo-dT and MMLV reverse transcriptase (Promega, M1705). The resulting cDNA pool was treated with 3 units of RNase H (Enzymatics, Y9220L) for 30 minutes at 37°C, and then analyzed by qPCR using a Roche LightCycler 480 system (initial 95°C for 5 minutes, 45 cycles of amplification at 95°C for 10 seconds, 60°C for 10 seconds, 72°C for 1 second) with SYBR Green detection and gene-specific primers (see list below). Target gene expression was normalized to the expression of RPL19 mRNA.

#### qPCR primer sequences

We used the following nucleic acid oligonucleotides for qPCR:

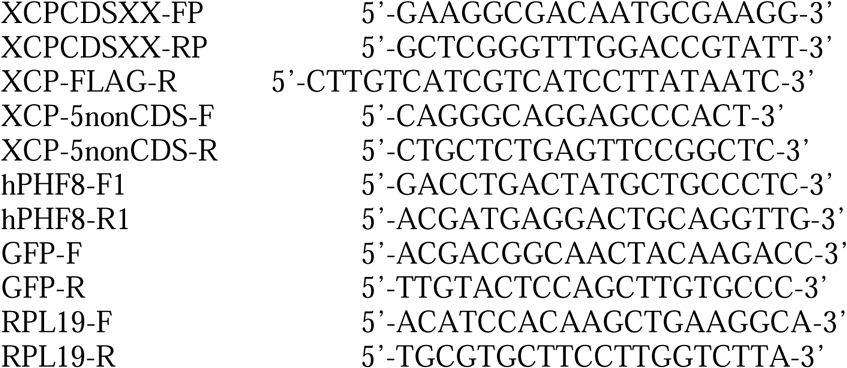

### Analysis of TCGA and GTEx Data

Data from TCGA and GTEx was downloaded from the recount2 database [43] that was aligned to the human reference genome (GRCh38/hg38) and gene annotations for reference chromosomes from GENCODE (v.25).

***Box Plots.*** Box plot representations were used to quantitatively assess the expression of genes in normal tissue. Additionally, box plot representations of the mean and standard error were used to assess the expression of genes in normal and cancer tissues. Differential expression of RPKM normalized data was tested by ANOVA, using the aov packaged in R.

### Histology and immunostaining

A breast disease spectrum tissue microarray was purchased from US Biomax (BR2082b). Immunohistochemical staining was performed on a Dako Autostainer Link 48 system. Briefly, 5 μm paraffin sections were baked for 20 minutes at 60^°^C, then deparaffinized and hydrated before the antigen retrieval step. Heat-induced antigen retrieval was performed at pH 6 (XCP) for 20 minutes in a Dako PT Link. The tissue was incubated with a peroxidase block and then an antibody incubation (1:100 XCP) for 20 minutes. The staining was visualized using the EnVision FLEX visualization system. The intensity of staining was scored on a scale of 0-3, where 3 is the highest intensity (expression).

### Generation of ectopic cell lines

#### Molecular cloning of XCP and PHF8

*LncRNA1456* 5’ and 3’ ends were mapped by RACE as described above. The full-length coding sequence was cloned by PCR with a 3’-FLAG epitope tag flanked by NheI and XhoI sites using MCF-7 cDNA. The full length *lncRNA1456* (wild-type and ATG mutant sequences), and PHF8 cDNAs were synthesized by Invitrogen GeneArt Gene Synthesis and sequence verified. Purified plasmid contained NheI and XhoI flanked full length *lncRNA14561* (wild-type or ATG mutant sequences), or PHF8 with 3’-FLAG epitope-tag coding sequence.

#### Lentiviral expression vector

The cDNAs, or a GFP-FLAG cDNA control, were inserted into pInducer20 lentiviral doxycycline (Dox)-inducible expression vector by ligation using NheI and XhoI sites.

#### Doxycycline-inducible expression in cell lines

Lentiviruses were generated by transfecting each pInducer20 vector into 293T cells, together with expression vectors for the VSV-G envelope protein (pCMV-VSV-G, Addgene plasmid no. 8454), the expression vector for GAG-Pol-Rev (psPAX2, Addgene plasmid no. 12260), and a vector to aid with translation initiation (pAdVAntage, Promega) using Lipofectamine 3000 transfection reagent (Thermo Fisher Scientific, L3000015) according to the manufacturer’s instructions. After 24 hours, culture medium was replaced with fresh medium and the cells were maintained for an additional 24 hours. The virus-containing supernatants were collected, filtered through a 0.45 μm syringe filter, and concentrated by using Lenti-X Concentrator (Clontech, 631231). The filtered supernatants were used to infect MCF-7 or MDA-MB-231 cells supplemented with 1 μg/mL polybrene (Sigma-Aldrich, H9268) to increase transduction efficiency. The infected cells were placed under selection with 1 mg/mL Geneticin/G418 (Thermo Fisher Scientific, 11811031), and once stable, maintained with 400 μg/mL Geneticin/G418. For doxycycline (Dox; Sigma-Aldrich, D9891) induction, cells were treated with 250 ng/mL of Dox for 24-48 hours. Expression of proteins was confirmed by Western blotting.

### Xenograft assays

All animal experiments were performed in compliance with the Institutional Animal Care and Use Committee (IACUC) at the UT Southwestern Medical Center. Female NOD *scid* gamma (NSG) mice at 6-8 weeks of age were used. To establish breast cancer xenografts, MDA-MB-231 or MCF-7 cells (5 x 10^6^ in 100 μl), engineered for doxycycline-inducible expression of FLAG-tagged XCP or GFP were injected subcutaneously into the flank of mice in a 1:1 ratio PBS and matrigel (Corning, 354234). For MCF-7 experiments, NSG mice were supplemented with 1.7 mg 17-β-estradiol 60-day release pellets (Innovative Research, SE-121) implanted subcutaneously at the base of the neck under anesthesia. Ten days post-tumor cell injection, mice were placed on a doxycycline containing diet (625 mg/kg; Envigo, TD.01306). Mouse weight was monitored once a week and tumor growth measured over time using electronic calipers approximately every 3-4 days. Tumor volumes were calculated using a modified ellipsoid formula: *Tumor volume* = ½ (*length* × *width*^2^). Animals were euthanized at 50 days post-injection. Fresh tumor samples were fixed for 24 hours in 10% formalin.

### Immunofluorescent staining and confocal microscopy

MCF-7 cells were seeded on four-well chambered cover slips (Thermo Fisher, 155411) 1 day prior to fixing. The following day, cells were washed three times with PBS, fixed in 3.7% paraformaldehyde for 15 minutes at room temperature, and washed with 0.1% PBS-Tween 20 (PBS-T). Cells were permeabilized for 5 minutes using 0.5% PBS-T, washed three times with 0.1% PBS-T, and incubated for 1 hour at room temperature in Blocking Solution (10% calf serum in PBS containing 0.5% gelatin) and then washed twice with 0.1% PBS-T. Fixed cells were incubated at room temperature with a polyclonal antibody against XCP or PHF8 in Incubation Solution (PBS containing 1% BSA) for 2 hours at room temperature and washed four times with 0.1% PBS-T. Then the fixed cells were probed with Alexa Fluor 488 goat anti-rabbit IgG (Thermo Fisher Scientific, R37116) in blocking solution for 1 hour, washed four times with 0.1% PBS-T, and incubated at room temperature with TO-PRO™-3 Iodide nuclear dye (Thermo Fisher Scientific, T3605) or 4,6-diamidino-2-phenylindole nuclear dye (Thermo Fisher Scientific, D1306) in PBS for 5 minutes, and then washed four times with 0.1% PBS-T again. Lastly, cells were treated with VectaShield (Vector Laboratories, H-1000) and imaged using a Zeiss LSM880 confocal microscope, purchased with a shared instrumentation grant from the NIH (1S10OD021684-01 to Katherine Luby-Phelps).

### In vitro pull-down assays

One microgram of the His_6_-tagged protein XCP was mixed with two micrograms of FLAG-PHF8 in the interaction buffer (20 mM Tris-HCl [pH 7.9], 20% glycerol, 0.2 mM EDTA [pH 8.0], 0.1% Nonidet P-40, 2 mM phenylmethylsulfonyl fluoride, 150 mM KCl, 30 mM imidazole) along with the Ni-nitrilotriacetic acid (NTA) His Bind resin (Qiagen). The mixture was incubated for 3 hours at 4°C on a rotary shaker. After the beads were extensively washed in the interaction buffer, the proteins were extracted from the beads into the sodium dodecyl sulfate (SDS) sample buffer, separated on an SDS–12% polyacrylamide gel electrophoresis (PAGE) gel, and visualized by Western blotting with rabbit anti-XCP and mouse anti-FLAG.

### Analysis of domain family

We downloaded the list of Homeobox family of proteins from the HGNC database [44]. All 314 Homeobox family proteins and the identified ORF were aligned using Clustal Omega [45].

### Analysis of *lncRNA1456*– and PHF8-regulated genes by RNA-seq

PolyA+ RNA-seq libraries were prepared from control and *lncRNA1456* knockdown MCF-7 cells as described above using the dUTP method [36]. Two biological replicates were generated for each sample. The RNA-seq raw reads were mapped to the hg19 human reference genome by TopHat [46], using RefSeq gene annotations as the reference for alignment. To determine expressed transcripts, we used custom R scripts to calculate the counts and RPKM using the Bioconductor packages GenomicRanges and edgeR (Lawerence et al., 2013, Robinson et al., 2010). Differentially regulated RefSeq mRNAs were called by Cuffdiff, using a 5% FDR, comparing the control samples to the lncRNA knockdown samples. We derived a high-confidence regulated mRNA set by filtering the Cuffdiff-called regulated mRNA lists with a fold cutoff of either 2^(0.8) or 2^(–0.8) for each siRNA-treated condition relative to the control. The resulting mRNAs with their corresponding fold changes were represented in heatmaps using Java Treeview. We also performed a transcription factor target analysis using Genomic Regions Enrichment of Annotations Tool (GREAT) [47] on the regulated mRNA set to draw inferences about transcription factors that may contribute to the observed gene regulation.

### Chromatin immunoprecipitation (ChIP)-sequencing and analysis

#### Chromatin immunoprecipitation

For ChIP assays, cells were cross-linked with 1% formaldehyde in PBS for 10 minutes at 37^°^C, quenched by the addition of 125 mM glycine, and incubated 5 minutes at 4^°^C. The cross-linked cells were collected in ice-cold PBS and pelleted by centrifugation. The cells were then resuspended by gentle mixing by pipetting in ice-cold Farnham Lysis Buffer (5 mM PIPES pH 8.0, 85 mM KCl, 0.5% NP-40) with freshly added 1 mM DTT and 1x protease inhibitor cocktail. The supernatant was removed, and the crude nuclear pellet was collected by centrifugation and resuspended in SDS Lysis Buffer (50 mM Tris-HCl pH 7.9, 1% SDS, 10 mM EDTA) with freshly added 1 mM DTT and 1x protease inhibitor cocktail. After a 10-minute incubation on ice, the lysate was sheared by sonication using a Bioruptor (Diagenode) to generate chromatin fragments of approximately 250 bp in length. The sheared chromatin was clarified by centrifugation and the diluted ten-fold in ChIP Dilution Buffer (20 mM Tris-HCl pH 7.9, 0.5% Triton X-100, 2 mM EDTA, 150 mM NaCl) with freshly added 1 mM DTT and 1x protease inhibitor cocktail. The lysate was pre-cleared with equilibrated protein A-agarose beads and subjected to immunoprecipitation reactions with an antibody against PHF8 (Bethyl, A301-772A) at 4^°^C overnight.

The immunoprecipitates were collected by incubation with BSA-blocked protein A-agarose beads for 2 hours at 4^°^C with gentle mixing. After incubation, the beads were washed on ice once each with (1) Low Salt Wash Buffer (20 mM Tris-HCl pH 7.9, 2 mM EDTA, 125 mM NaCl, 0.05% SDS, 1% Triton X-100, 1x protease inhibitor cocktail), (2) High Salt Wash Buffer (20 mM Tris-HCl pH 7.9, 2 mM EDTA, 500 mM NaCl, 0.05% SDS, 1% Triton X-100, 1x protease inhibitor cocktail), (3) LiCl Wash Buffer (10 mM Tris-HCl pH 7.9, 1 mM EDTA, 250 mM LiCl, 1% NP-40, 1% sodium deoxycholate, 1x protease inhibitor cocktail), and (4) 1x Tris-EDTA (TE). The beads were then subjected to a final wash with 1x TE at room temperature. They were then collected by centrifugation, resuspended in ChIP Elution Buffer (100 mM NaHCO3, 1% SDS), and incubated on end-over-end rotator for 15 minutes at room temperature to elute the ChIPed DNA. The ChIPed DNA was de-crosslinked by adding 100 mM NaCl with incubation at 65^°^C overnight. The eluted material was cleared of protein and RNA by adding RNase H and proteinase K, and incubating for 2 hours at 55^°^C. The ChIPed DNA was then extracted with phenol:chloroform:isoamyl alcohol (25:24:1), collected by ethanol precipitation, and dissolved in water. ChIPed DNA was used for library preparation for sequencing.

#### Library preparation

ChIP libraries were prepared using a modified KAPA LTP Library Preparation kit (KAPA BiosystemsKK8232) for Illumina Platforms. Ten ng of sheared DNA was used to repair the ends of the damaged fragments using a proprietary master mix. The resulted blunted fragments were 3′ A-tailed using a proprietary mixture of enzymes to allow ligation to the specific Illumina adaptors. Each of the steps (i.e., end repair, 3’A tailing, and adaptor ligation) was followed by AMPure XP bead clean up (Beckman Coulter, A63881). After adapter ligation, DNA enrichment was performed using Kapa HiFi Hot Start Ready PCR mix, and a cocktail of primers (1 cycle at 98 °C for 45 seconds; 5 cycles at 98 °C for 20 seconds, 63 °C for 30 seconds, and 72 °C for 30 seconds; and 1 cycle at 72 °C for 1 min), and purified with AMPure XP beads. DNA templates were size selected (∼200-300 bp) by running on a 2% agarose gel, followed by PCR enrichment (1 cycle at 98 °C for 45 seconds; 11 cycles at 98 °C for 20 seconds, 63 °C for 30 seconds, and 72 °C for 30 seconds; and 1 cycle at 72 °C for 1 min) and final purification. The quality of the final libraries was assessed using a 2200 TapeStation (Agilent Technologies). The libraries were quantified using Qubit dsDNA High Sensitivity Assay Kit (Thermo Fisher Scientific, Q32854) and samples pooled at final concentration of 2 nM.

#### Library sequencing

The libraries were sequenced using a NextSeq sequencer (Illumina; Single-end reads, 75 bp for all samples). At least two biological replicates were sequenced for each cell line for a minimum of roughly 100 million raw reads per cell line.

#### Quality control

Quality control for the RNA-seq data was performed using the FastQC tool (http://www.bioinformatics.babraham.ac.uk/projects/fastqc/).

#### Read alignment and peak calling

The raw reads were aligned to the human reference genome (GRCh37/hg19) using default parameters in Bowtie (ver. 1.0.0) [48]. The aligned reads were subsequently filtered for quality and uniquely mappable reads using Samtools (ver. 0.1.19) [49] and Picard (ver. 1.127; http://broadinstitute.github.io/picard/). Library complexity was measured using BEDTools (v2.17.0) [38] and met minimum ENCODE data quality standards [50]. Relaxed peaks were called using MACS (v2.1.0 [51] with a p-value = 1 x 10-2 for each replicate, pooled replicates’ reads, and pseudoreplicates. Called peaks from the pooled replicate that were observed in both replicates or in both pseudoreplicates were used for subsequent analyses.

#### Metagene and heatmap analysis

The average read densities of reads were calculated for an 20kb window surrounding peak center (±10 kb) using the plotProfile and plotHeatmap functions in deeptools (ver. 2.5.0.1) [52].

### Demethylation assay and dot blots

Purified His_6_-XCP, recombinant Histone H3K9me2 (Active Motif, 31280), and recombinant FLAG-PHF8 protein (Active Motif, 31435) were used for in vitro assays. Ten microliters of each demethylation reaction were spotted onto a nitrocellulose membrane. The membranes were air dried, blocked with 5% nonfat milk in TBST, incubated with the Histone H3K9me2 (Cell Signaling, 9753) or Histone H3 (Abcam, ab1791) primary antibodies described above in 3% non-fat milk prepared in TBS-T, and then incubated with goat anti-rabbit HRP-conjugated IgG (Pierce, 31460). Western blot signals were detected using an ECL detection reagent (Thermo Fisher Scientific, 34077, 34095).

### Histone post-translational modification mass spectrometry and analyses

Histone analysis was carried out as previously described [53]. After isolation of nuclei, acid extraction was performed to release histone proteins from nuclei. Chemical derivatization was carried out for mass spectrometry analysis of histone modifications. The extracted histone sample was dissolved in ammonium bicarbonate, and its pH was adjusted to 8. Propionylation reagents mixing propionic anhydride and acetonitrile (ACN) in a 1:3 ratio (v/v) were added, followed by NH4OH treatment to regulate pH levels. Centrifugation and vacuum drying produced propionylated histones, ideal for proteolytic digestion and HPLC-mass spectrometry analysis. Additionally, Western blotting was performed to analyze histone modifications. Histones extracted from cultured cells were treated with SDS-PAGE Loading Buffer and heated. Western blotting was conducted on a 15% SDS-PAGE gel, with ∼10 µg of histone proteins, and histone modifications were detected using specific antibodies.

The Mass-spectrometry spectra were analyzed using Proteome Discoverer v2.4 SP1 (Thermo Fisher Scientific, Waltham, MA). To mine this data for modifications of interest, we first filtered the records in the mass spectrometry output file to contain only those which were indicated to be contained within either the Histone H3 or H4 complex, of these we then determined which entries covered the sites of interest (H3K9). After determining a listing of all records in the mass spectrometry data that covered these sites, we calculated the relative abundances of PTM p at that site (k) for each replicate (t) of each condition (c) via:

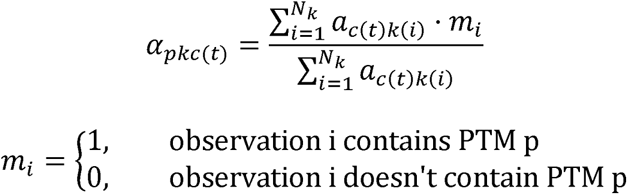

Where α*_pkc_*_(*t*)_ is the relative abundance of PTM p at site k, of replicate t of condition c, α*_c(t)k(i)_* is the i^th^ reported absolute abundance of replicate t of condition c at site k, *N_k_* is the total number of records covering site k, and *m_i_* is an indicator of whether or not a modification p is present in observation i of site k. Software, scripts and other information about the analyses can be obtained by contacting the corresponding author (W.L.K.).

### Data availability

All of the RNA-seq, ChIP-seq and mass spectrometry data sets used in this study can be accessed through the NCBI’s Gene Expression Omnibus (GEO) repository (http://www.ncbi.nlm.nih.gov/geo/) using accession number GSE288868 for RNA-seq, and GSE287423 for ChIP-seq, or the Mass Spectrometry Interactive Virtual Environment (MassIVE) repository (https://massive.ucsd.edu/) using the accession number MSV000096645.

