## Supplementary Data for "X-Linked Cancer-Associated Polypeptide (XCP) from *lncRNA1456* Cooperates with PHF8 to Regulate Gene Expression and Cellular Pathways in Breast Cancer"

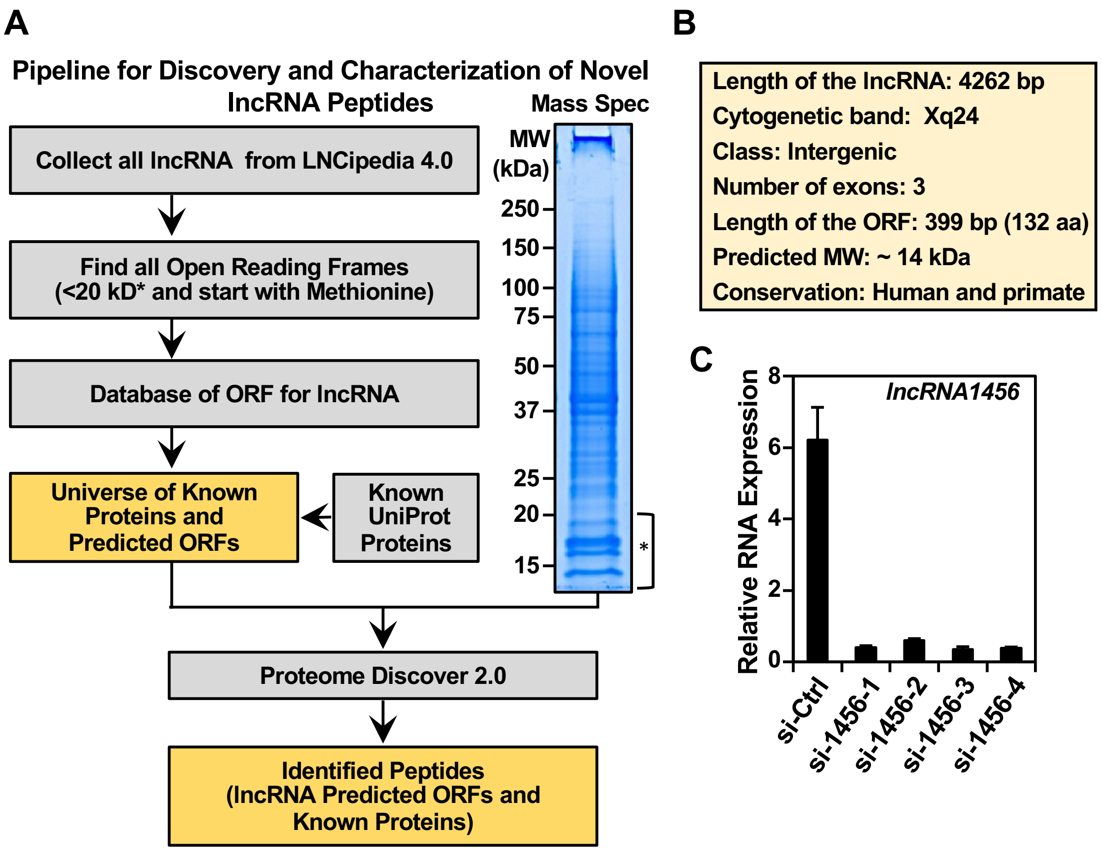


Suppl. Fig. S1 – Gadad *et al.* (2025)

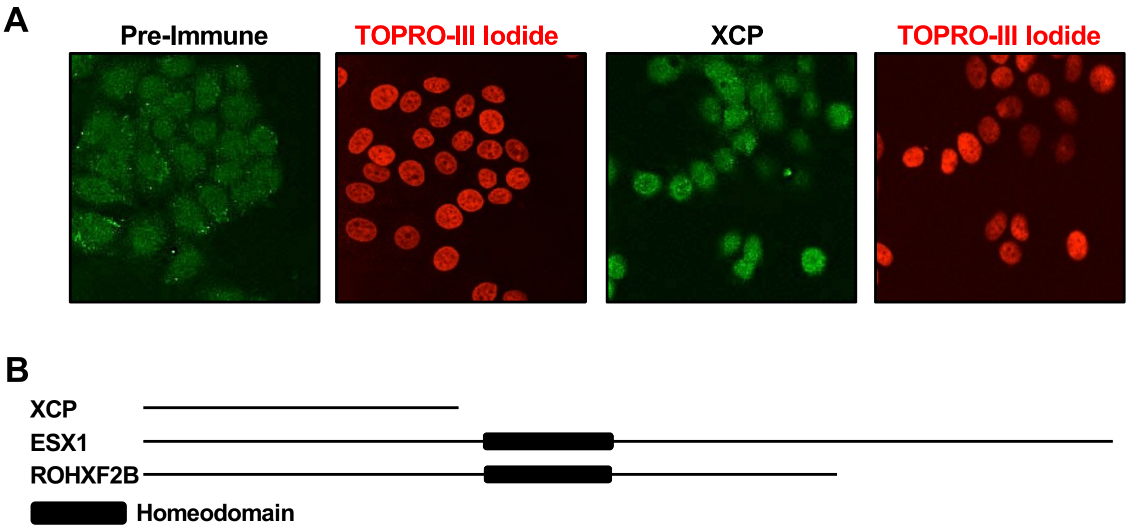


Suppl. Fig. S2 – Gadad *et al.* (2025)

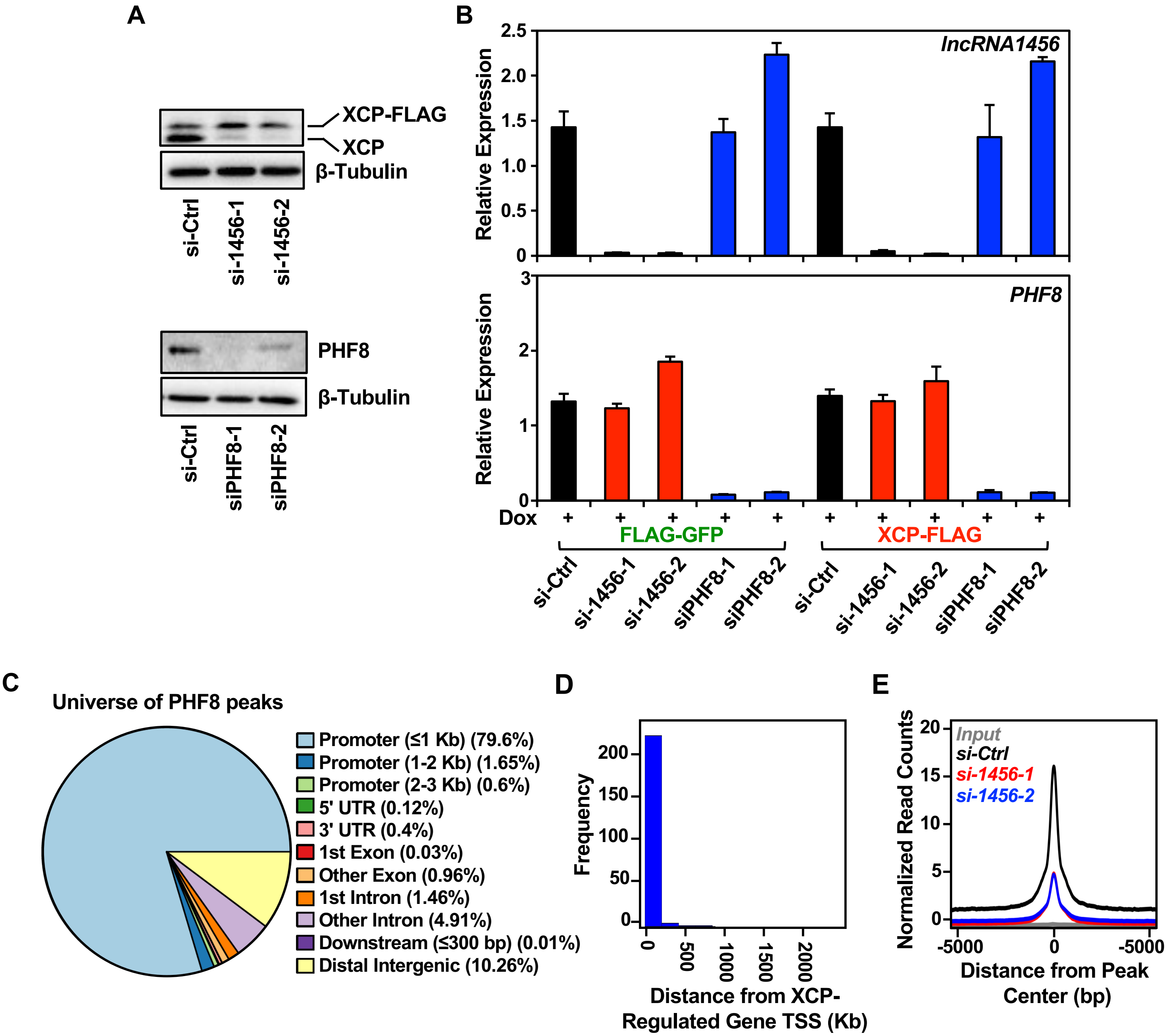


Suppl. Fig. S3 – Gadad *et al.* (2025)

**Table S1. PH8 Identification by Mass Spectrometry Analysis**

| Protein | Length (AA) | mw (KDa) | PSMs | Peptide Seqs | % Coverage | XCP-150/ GFP-150 | XCP-200/ GFP-200 |
|---|---|---|---|---|---|---|---|
| EGFP | 248 | 28.122 | 973 | 42 | 91.5 | 0.01 | 0.01 |
| XCP | 132 | 13.753 | 363 | 12 | 90.9 | 595.04 | 271.67 |
| PHF8 | 953 | 114.148 | 5 | 5 | 8 | 11.48 | 6.62 |
| H3 | 136 | 154.56 | 2 | 6 | 35.3 | XCP-150 only | XCP-200 only |
